# Combining prior knowledge and transcriptomics data for logic models of patient subgroups

**DOI:** 10.64898/2026.06.19.732519

**Authors:** Bi-rong Wang, Yunfan Bai, Julio Saez-Rodriguez, Federica Eduati, Aurélien Dugourd

## Abstract

Computational modeling provides a powerful framework for in silico exploration of anti-cancer therapeutic targets and tumor response mechanisms. Oncogenic signaling pathways play a central role in tumor behavior and represent promising targets for personalized combination therapies. However, these pathways are complex, and although logic-based models are well suited for representing signaling dynamics, they are often constrained by model-specific data requirements, limited scalability, and time-consuming manual curation. Here, we introduce Functional Integration of Contextualized Omics for Unraveling regulatory dynamicS (FICUS), a framework that integrates omics-driven network contextualization with dynamic Boolean and logic-ODE modeling. FICUS enables automated, data-driven protein network inference and patient stratification, allowing shared signaling mechanisms to be identified across patient subgroups while preserving patient-specific dynamic responses. We applied FICUS to the SU2C-MARK lung cancer cohort and the The Cancer Genome Atlas kidney cancer cohort, demonstrating its utility for post-hoc analyses and downstream interrogation of dynamic tumor models. Overall, our results highlight the flexibility of FICUS in capturing heterogeneous signaling mechanisms across patient subgroups, addressing a key challenge in precision oncology.

## Introduction

Tumor heterogeneity, across patients, lesions or even individual cells, remains a major challenge in precision oncology. Addressing this complexity is essential for improving long-term efficacy of solid tumor therapies and preventing drug resistance, metastasis, and recurrence[1]. Overcoming resistance requires a detailed understanding of patient-specific responses, enabling development of personalized treatment strategies tailored to individual molecular profiles. However, empirically testing therapy combinations for each patient is time-consuming, burdensome, and often associated with significant toxicities. However, if therapy combinations are successfully found, they can develop into valuable strategies for treating highly malignant and aggressive cancers such as pancreatic cancer [2]. In this context, computational modeling offers a promising framework to screen for effective therapies, allowing in silico exploration of therapeutic targets and the mechanisms underlying tumor responses. In particular, oncogenic signaling pathways, though complex, are central to tumor behavior and provide a rich source of targets for personalized combination therapies[3].

High-throughput omics technologies have enabled systematic investigation of signaling pathways at multiple biological levels, including genomics, transcriptomics, proteomics, and metabolomics. Numerous computational tools have been developed to analyze such data and extract mechanistic insights[4–6]. Increasingly, these data are integrated into multi-omics datasets to capture complementary regulatory layers. For example, TieDIE was used to integrate multi-omics data for molecular characterization of clear cell renal cell carcinoma [7], while COSMOS (Causal Oriented Search of Multi-Omics Space) links signaling and metabolic networks through transcription factor, kinase/phosphatase, and metabolite activities inferred from omics data [8,9]. Such approaches enable scalable, data-driven inference of context-specific signaling networks. However, the resulting representations are inherently static, limiting their ability to simulate dynamic tumor responses or treatment trajectories [10].

Dynamic logic models offer a complementary framework well suited for simulating signaling pathway responses and generating mechanistic hypotheses. Due to their interpretability and relatively low computational complexity, logic-based models scale well to large signaling networks. Although logic models are relatively coarse, limiting their description of biochemical details, such formalism allows logic models to not be dependent on complex molecular information. Instead, prior knowledge on for instance signaling networks can be incorporated to compensate for the limited molecular detail. Montagud et al. (2022), for example, developed personalized Boolean logic models of deregulated signaling pathways in prostate cancer, constrained with baseline patient omics data (i.e. mutations, CNVs and gene expression)[11]. Eduati et al. (2020) further demonstrated how CellNOpt can create logic-derived ordinary differential equation (ODE) models of apoptosis mechanisms, optimized with patient-specific perturbation data[12]. By combining OR- and AND-gates, such models capture biologically meaningful regulatory logic, which can represent e.g. cooperative activation and protein complex formation.

Despite their promise, the widespread application of mechanistic logic models is constrained by their dependency on data, e.g. protein activity readouts, specific to the model structure. For instance, applications of the logic-ODEs from Eduati et al. (2020) require a snapshot of multiple perturbations per sample (e.g. cell line, patient) to enable model fitting and ensure identifiability[12]. While such perturbation experiments can be systematically performed at large scale in cell lines, they are generally not feasible in patients. Consequently, patient-derived omics datasets, whether generated by sequencing, mass spectrometry, or antibody-based approaches, typically consist of a single baseline measurement per patient. As a result, although large-scale omics data are available for patient cohorts and have been used in several instances to support the development of logic and dynamic models [11,13], the lack of perturbation conditions limits their direct use for fitting dynamic logic-ODE models, restricting their applicability in clinical settings. A second major limitation concerns model scalability and construction. While approaches such as Montagud et al. (2022) successfully leverage omics data for model personalization, the underlying signaling networks still require extensive manual curation, constraining adaptability and scalability [11,14]. Similarly, CellNOpt relies on prior knowledge networks (PKNs), and although these can be created through both manual or data-driven curation, in practice they are typically assembled through manual curation process [15]. While tools such as NeKo exist to generate logic models automatically from multiple databases, they do not leverage quantitative measurements to constrain activity flows or build patient-specific networks [16]. To date, no scalable framework simultaneously leverages omics data to define model structure and inform dynamic models at both group and patient levels. In this context, omics-derived static networks provide a promising starting point for automated model contextualization.

Based on the potential complementarity between omics-derived static networks and dynamic logic modeling, we introduce Functional Integration of Contextualized Omics for Unraveling regulatory dynamicS (FICUS). This framework integrates omics-derived network contextualization with dynamic Boolean and logic-ODE modeling. Instead of constructing PKNs bottom-up from literature, FICUS employs COSMOS to estimate protein activities and contextualize signaling networks in a top-down manner using patient omics data. To address the lack of perturbation data in patient cohorts, we further leverage inter-patient variability as a source of information for model training. This strategy enables the inference of common mechanisms while generating patient-specific dynamic responses, thereby supporting subgroup analysis and the investigation of heterogeneous tumor behavior, which is a key challenge in precision oncology [17].

We used the human prostate cancer patients from TCGA to test the automated curation of the PKN [11,18]. Next, we demonstrated the application of FICUS using a lung cancer transcriptomics cohort and the TCGA KIRC dataset. Our results show that automated network contextualization at both individual and subgroup levels yields dynamic models capable of capturing shared and patient-specific regulatory mechanisms. Finally, we present multiple strategies for model analysis and *in silico* perturbation experiments to generate mechanistic hypotheses, providing a foundation for future experimental and therapeutic exploration.

## Results

### Setting up fitting models to patient subgroups

We developed a multi-step workflow (FICUS) to derive patient-specific signaling models from cohort omics data (**Figure 1**). Starting from transcriptomics data, we first inferred transcription factor activities and curated patient-specific protein networks using COSMOS, followed by functional scoring. These outputs were then converted into CellNOpt-compatible inputs to enable logic model optimization. Unlike conventional applications of CellNOpt, which rely on perturbation response data to generate models for individual samples or cell lines (Eduati et al. 2020), our approach leverages transcriptomics data with only a single measurement per patient. To address the resulting limitation of insufficient data per individual, we exploited inter-patient heterogeneity as a source of variation to inform model optimization. Using this strategy, we constructed models that capture mechanisms shared within patient subgroups while preserving patient-specific features. Overall, FICUS comprises data preprocessing, COSMOS-based network inference, transformation into CellNOpt inputs, model optimization, and downstream analysis of the resulting logic models.

**Figure 1:**
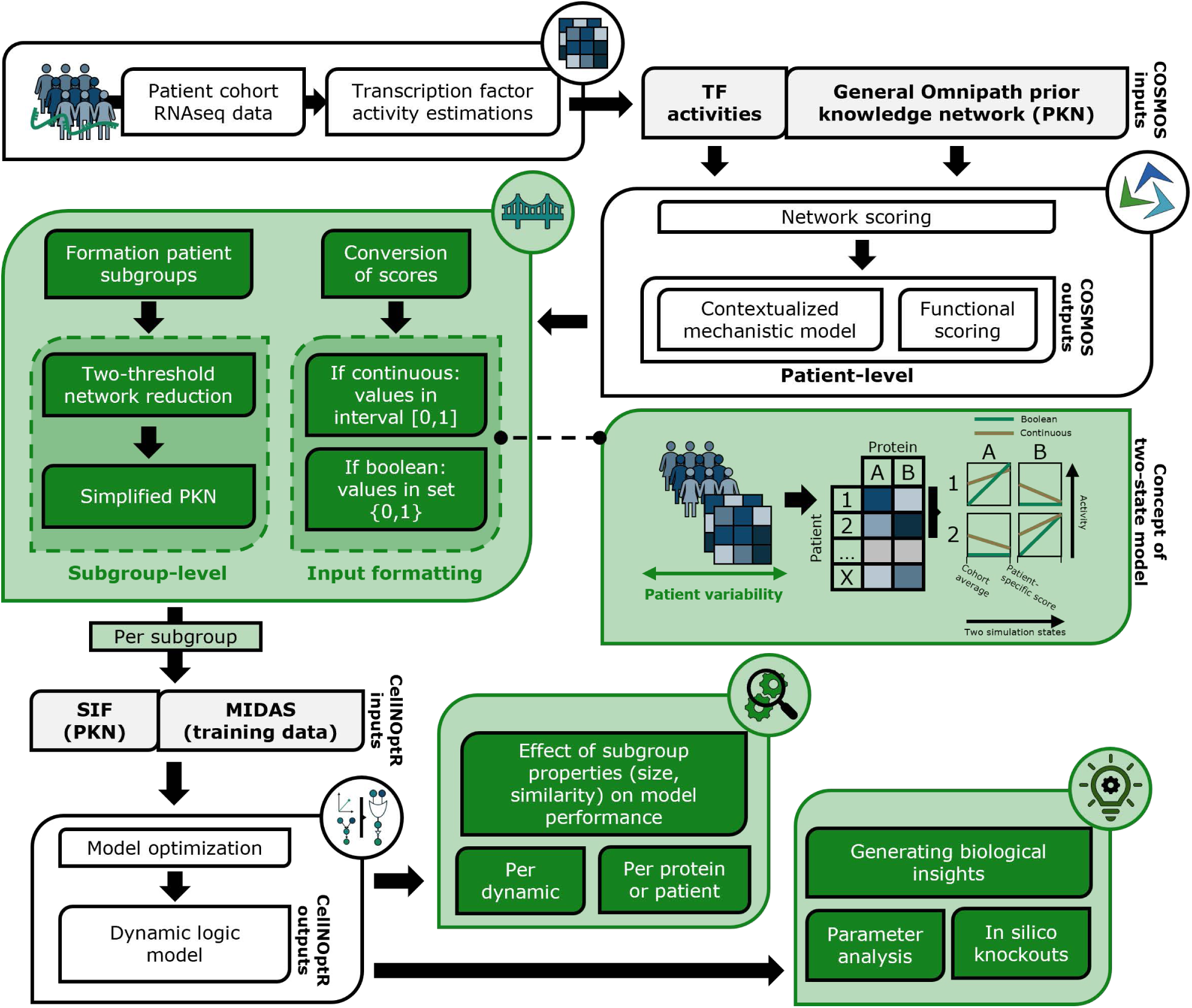
Overview FICUS. An overview of the steps in FICUS, using omics data to build and inform logic models. Blocks in green highlight the new implementations to support the link between data and tools such as COSMOS and CellNOpt.

We applied two logic formalisms from CellNOpt to construct patient subgroup models: Boolean and logic-ODE [19]. The Boolean model only produces binary outputs (0 or 1), while the logic-ODE model generates continuous values between 0 and 1. Even though continuous models are expected to better represent biological complexity, Boolean models can be optimized much faster, which was valuable for setting up our approach. Both formalisms require a prior knowledge network (PKN) and corresponding training data, which we derived from patient omics profiles using MOON, a component of COSMOS [9]. MOON estimated functional scores for proteins and outputs patient-specific networks from a generic Omnipath PKN [20]. For more details related to MOON, see **Methods**. We then converted functional scores into training data for the models and aggregated patient-specific networks into subgroup-specific PKNs. Through this process, we generated optimized dynamic logic models for patient subgroups, enabling us to capture both shared and context-dependent mechanisms across the cohort.

### Curating prior knowledge networks of patient subgroups through clustering and network reduction

Excessive heterogeneity can obscure shared mechanisms and increase the presence of contradicting signaling features, which are common in cancer. Therefore, to address patient heterogeneity, we divided the cohort into subgroups rather than fitting a single model to all patients. Patient-specific protein networks were first inferred from transcriptomics data, representing candidate sets of proteins and interactions capable of explaining the observed transcription factor activities in each patient. We then grouped patients based on the similarity of these curated networks, with the aim of identifying recurrent regulatory structures across patients. In this context, we interpret such recurring network structures as shared mechanisms. We generated subgroups using hierarchical clustering and we also randomly generated subgroups to serve as a baseline to evaluate the effect of clustering. The dendrogram cut used to define the number of clusters was determined for each data set separately (see **Methods** for more details). For hierarchical clustering, we measured pairwise similarity between patient-specific networks using the Jaccard index, thereby grouping patients with similar regulatory structures. This approach enables the identification of subgroups that capture common regulatory dependencies (i.e. shared mechanisms) while retaining sufficient variability to support model optimization.

For each subgroup, we aggregated the patient-specific networks into a combined PKN for logic model optimization (union across networks). These aggregated networks initially contained thousands of interactions, which posed challenges for optimization, particularly in the logic-ODE formalism. To reduce network size, we implemented a double-threshold reduction strategy (see **Methods**) that selected top proteins based on average functional scores while retaining direct neighbors to preserve connectivity. Increasing thresholds leads to stronger reductions, which if desired can reduce PKNs to have less than e.g. 50 interactions.

We further refined the PKNs by node and edge filtering to reduce sparsity in training data and minimizing incoherent protein interactions across patients. Together, these steps reduced network complexity while retaining subgroup consensus, resulting in combined PKNs that remained sufficiently large to capture relevant signaling mechanisms.

### Patient heterogeneity is integrated in the training data through patient-specific inputs and two simulation states

In perturbation-based models, different conditions are characterized by specific perturbations of the same biological system (e.g. a cell line), to capture different states of the underlying biological network. In our method, patients are abstracted as different states of a common underlying biological network (under the assumption that at least a part of their specific wiring might overlap), and therefore represent conditions in CellNOpt. However, the perturbed nodes of a given patient/condition are not a-priori known, differently from the classic perturbation experimental setting. To bypass this issue we defined a procedure to assign input nodes for each patient. For each patient-specific network, proteins without upstream interactions (“top nodes”) and significant activity scores were designated as stimuli (positive activity scores) or inhibitors (negative scores).

Functional scores were then converted into CellNOpt-compatible training data. For Boolean models, scores were discretized into stable (0) or upregulated (1) states relative to the cohort baseline. For logic-ODE models, scores were scaled to the [0,1] interval, with an initial baseline of 0.5 and final values reflecting patient-specific activities (**Figure 1**). These transformations enabled consistent integration of MOON outputs into both model formalisms.

### Curating patient group-specific protein networks from omics data

We applied the first part of FICUS (MOON and network clustering) to the TCGA prostate cancer data set [18] to infer patient-specific PKNs in a data-driven manner. The same dataset was previously used by Montagud et al. (2022) to personalize prostate cancer Boolean models based on a manually curated PKN [11]. Here, we aimed to assess whether automated network inference from omics data can overcome the bottleneck of manual PKN curation by comparing our inferred networks with the curated PKN from Montagud et al., hereafter referred as the Montagud_PKN.

The overlap between each patient-specific network and the Montagud_PKN varied per patient subgroup (**Figure 2A**). Subgroups 1-4 showed, on average, a higher overlap fraction (0.5-0.6) compared with subgroups 5-10 (0.1-0.4). Interestingly, while the overlap distributions within subgroups were relatively narrow, patient similarity within clusters varied. Pairwise Jaccard indices indicated that patients in subgroups 1-4 shared more interactions with each other (approximately 0.6-0.8) than patients in subgroups 5-10 (below 0.4; **Supplementary Figure 1**). Together, these results indicate that the Montagud_PKN shares more interactions with patient-specific networks from subgroups 1-4 than with those from subgroups 5–10, and that this overlap differs across subgroups.

**Figure 2:**
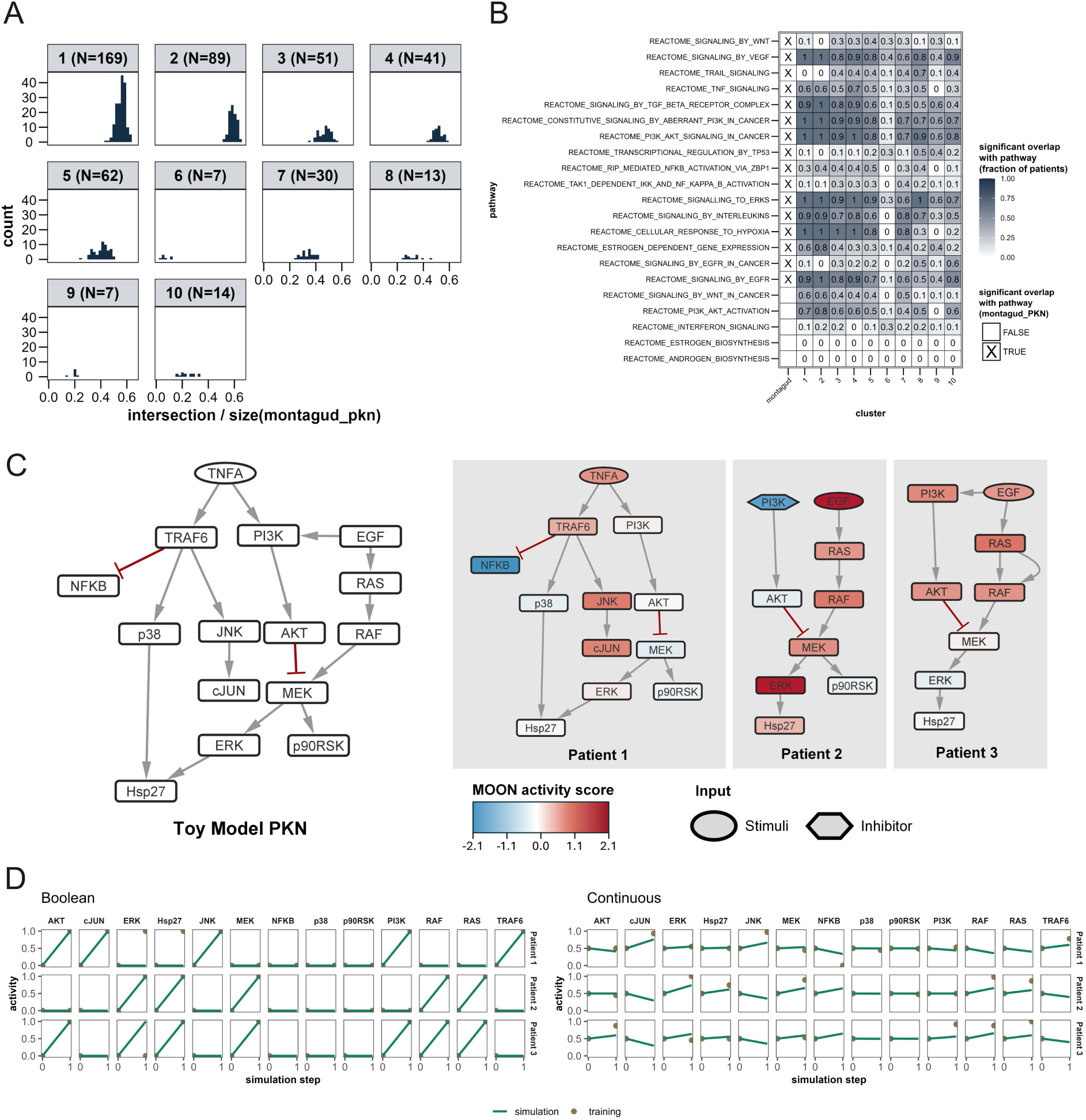
Automated protein network curation and conversion to CellNOpt inputs. **(A)** Comparison patient-specific protein networks with the prior knowledge network from Montagud et al. (2022), categorized per cluster the patient belongs to. The intersection (overlap in proteins between patient-specific networks and Montagud PKN) is normalized by the number of proteins in the Montagud PKN. The number of patients (N) is indicated per cluster. **(B)** Comparison overlap with gene sets across clusters and the Montagud PKN. For the patient clusters, a fraction is provided of how many patient-specific protein networks had significant overlap with the gene set of a pathway. For the montagud_PKN, the proteins from the network had either a significant or non-significant overlap with the pathways. **(C)** Individual protein networks of the three patients in the toy example, including the type of inputs per patient (stimuli or inhibitor) and the synthetically generated MOON activity scores for the different proteins. The combined PKN based on the three patients is given on the right as ‘Toy Model PKN’. **(D)** Model simulations after model fitting to inputs prepared from the toy example. Both Boolean and continuous (logic-ODE) formalisms were used to fit protein activities simultaneously for three patients.

To assess functional similarity, we evaluated the significance of overlap between the patient-specific protein networks per cluster and the Montagud_PKN with established pathway signatures using Fisher’s exact test. Pathway signatures were selected from Reactome gene sets matched to PROGENy pathways [21]. The Montagud_PKN showed significant overlap with most pathways, with the exception of SIGNALING_BY_WNT_IN_CANCER, PI3K_AKT_ACTIVATION, INTERFERON_SIGNALING, ESTROGEN_BIOSYNTHESIS, and ANDROGEN_BIOSYNTHESIS (**Figure 2B**). Notably, overlap patterns varied across clusters. For example, a larger fraction of cluster 2 networks showed significant overlap with PI3K_AKT_ACTIVATION (0.8) compared with clusters 5 (0.5) and 7 (0.4). Similarly, clusters 8 (0.5) and 10 (0.6) showed a higher proportion of significant overlap with SIGNALING_BY_EGFR than other clusters (≤0.4). Conversely, pathways with significant overlap in the Montagud_PKN were not always similarly represented across patient subgroups, such as SIGNALING_BY_WNT and TAK1_DEPENDENT_IKK_AND_NF_KAPPA_B_ACTIVATION.

Overall, while the Montagud_PKN shows significant overlap with a broad range of biological pathways, the extent of this overlap varies across patient subgroups. Several pathways captured by the Montagud_PKN are not uniformly represented in the patient-specific networks, indicating subgroup-specific differences in pathway involvement. These findings suggest that a single curated PKN may not fully reflect the heterogeneity of signaling processes across patient subgroups. Our approach in inferring patient-specific networks in a data-driven manner enables automated PKN inference, where the combination of MOON and network clustering could then capture both shared and context-specific biology without the need for manual curation.

### Toy models demonstrate how logic models can be optimized to MOON outputs

We tested our conversion from MOON outputs to CellNOpt using a toy model from CellNOpt [19]. To create a test case, we modified the stimulatory TRAF6(1)NFKB interaction into an inhibiting TRAF6(−1)NFKB interaction. This change introduced a protein with only inhibitory upstream interactions. The adjustment increased the tendency of the model to assign an initial activity value of 1 to NFKB. We created three synthetic MOON networks with functional scores (**Figure 2C**) corresponding to three partially overlapping subsets of a larger PKN. The protein networks were variations of the toy model from CellNopt, while MOON scores were randomly assigned to each protein. For both binary and continuous models, our approach successfully converted the patient-specific inputs into training data and a combined PKN suitable for model optimization. Using our ‘top nodes’ approach, key inputs such as TNFA, PI3K and EGF effectively distinguished the three patients. Model fitting captured patient-specific differences: the Boolean formalism reproduced distinct protein activities for each patient, while the logic-ODE formalism captured general trends. For example, MEK and PI3K activities varied between simulated patients as expected (**Figure 2D**). These results illustrate how our approach could convert omics data into CellNOpt-compatible inputs and generate logic models capable of simulating patient-specific responses while fitted to a group.

### Enrichment analysis of patient subgroups highlight differences between hierarchical clustering and random subgroups

Next, we applied our workflow to the SU2C-MARK lung cancer patient cohort [22], deriving patient-specific MOON outputs and clustering patients into subgroups. We analyzed the effect of defining patient subgroups through hierarchical clustering and compared the results with a baseline from randomly generated subgroups. Even though Jaccard indices are defined based on the network topology, we hypothesized that these differences might translate into distinct biological functions. The link to the biological context would be a valuable advantage of using hierarchical clustering. We therefore investigated biological functions of patient subgroups through an enrichment analysis using the hallmark gene set collection of MSigDB [23](See **Methods; Figure 3A**).

**Figure 3:**
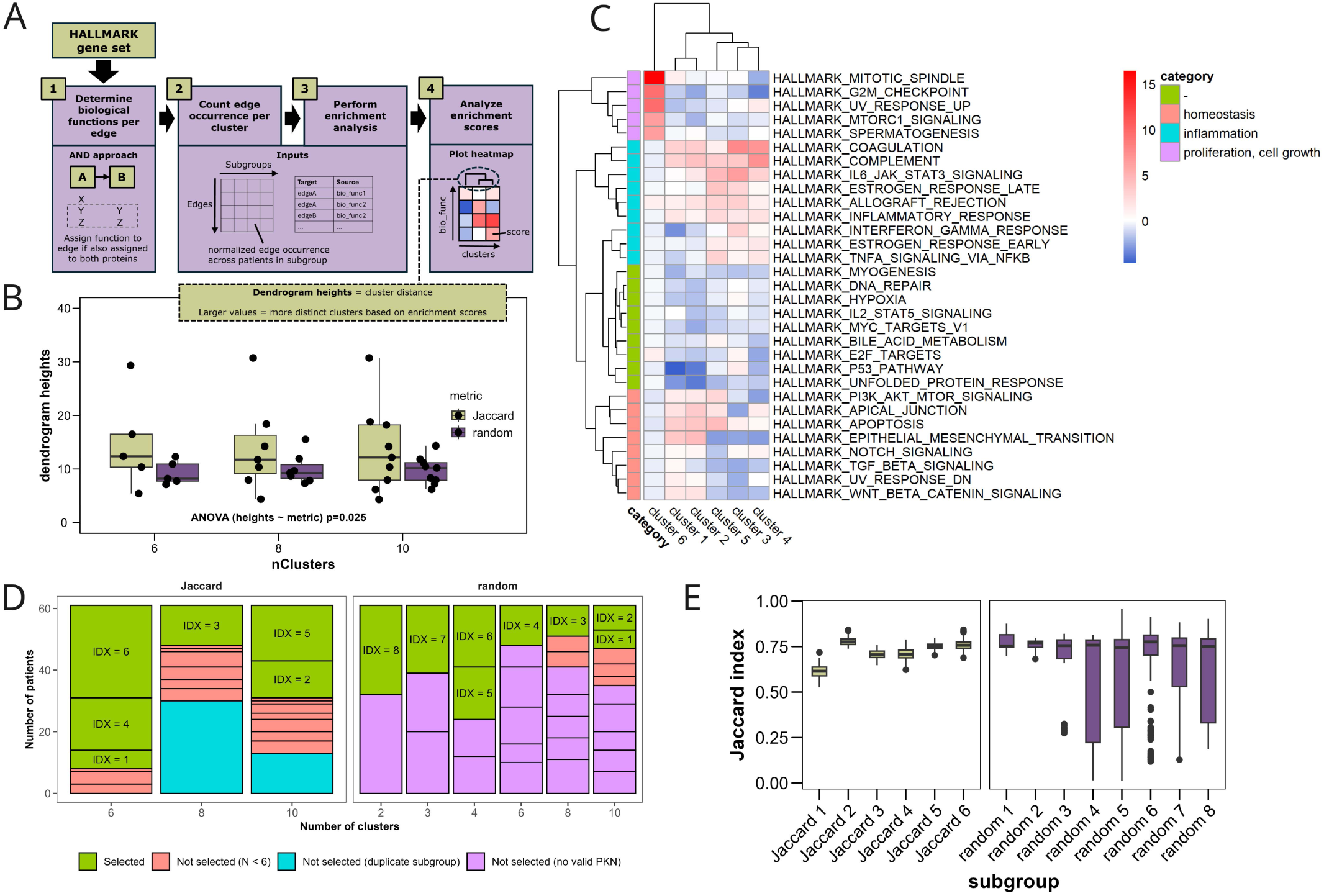
Cluster analysis of patient subgroups. **(A)**. Pipeline for the enrichment analysis of patient subgroups using the patients-specific protein networks. Instead of using biological functions per protein, as network edges were used for defining patient subgroups, this pipeline integrates the frequencies of edges and linked biological functions to derive enrichment scores. **(B)**. Comparison of dendrogram heights for different numbers of clusters derived from the hierarchical clustering and random subgroups. The dendrogram heights represent distances between clusters, i.e. bigger distances correlates with more distinction across clusters. Every data point represents a distance between two clusters. **(C)**. Enrichment scores per HALLMARK gene set and per cluster. Gene sets are also linked to general categories, e.g. homeostasis and inflammation. Enrichment scores can be positive (upregulated) or negative (downregulated). **(D)** Illustration of the selection heuristic for subgroups originating from different splits of the cohort. Jaccard subgroups originate from hierarchical clustering with Jaccard index as similarity metric across patient-specific networks. Selected subgroups (green) are additionally labelled with an IDX for easy reference. Unlabeled subgroups are colored based on the criteria that resulted in the non-selection (subgroup did not meet size criteria, was a duplicate or did not result in a valid PKN for the subgroup). **(E)** Comparison of all pairwise Jaccard indices across patients in a subgroup. Indices for Jaccard and random subgroups align with the IDX number visualized in **(D)**.

We analyzed the distribution of enrichment scores and dendrogram heights to visually represent distances between scores of each subgroup (**Figure 3A**). Jaccard subgroups exhibited greater variation in dendrogram heights compared to random subgroups (ANOVA, p=0.025; **Figure 3B**), indicating more distinct biological function enrichments. However, the median dendrogram heights decreased if the number of clusters (nClusters) created with hierarchical clustering was increased (4.17, 2.48, 1.96 for nClusters = 6, 8, and 10), consistent with the expectation that larger numbers of clusters result in less distinct clusters. Focusing on nClusters = 6, distinct biological functions were enriched in different Jaccard subgroups (**Figure 3C**). For example, cluster 6 showed strong upregulation in cell proliferation and growth, whereas inflammatory response pathways were more enriched in cluster 3. Clusters 3–5 exhibited stronger downregulation in pathways related to homeostasis and apoptosis. These results indicate that subgroups defined purely by network topology can also reflect meaningful biological distinctions.

### Cluster selection for investigating effect of subgroup properties

Selecting appropriate clusters for model optimization required balancing subgroup size and heterogeneity. Subgroups smaller than five patients risked overfitting, while very large subgroups could be too complex to capture patient-specific responses. Setting nClusters to 6, 8, or 10 often produced one or two large Jaccard subgroups (∼30 patients), with remaining clusters either below the minimum size or no common protein network could be generated. This latter issue, commonly found with random subgroups, arises due to patients having little to no overlapping proteins or edges, or due to network reduction and filtering dropping most of the network due to incoherence. We therefore decided to use subgroups clustered with different nClusters to impose variation in cluster size and increase the number of clusters to use for model optimization, resulting in six Jaccard subgroups and eight random subgroups (**Figure 3D**). Due to our selection heuristic to prioritize variation in subgroup properties, some subgroups have overlap in patients. For example, Jaccard subgroups IDX=2 (N=12) and IDX=5 (N=18) were a split of the IDX=6 (N=30) cluster, while IDX=3 (N=13) was a subset of IDX=4 (N=17) (**Supplementary Figure 2**).

Despite these overlaps, the selected subgroups provided sufficient variation to investigate the effects of cluster size on model performance. Sizes for both Jaccard and random subgroups ranged from 6 to 30 patients. Additionally, a comparison between subgroup similarities highlights an interesting difference between Jaccard and random subgroups: Jaccard subgroups have a reduced variation in pairwise Jaccard indices compared to random subgroups (**Figure 3E**). These results suggest that in the random subgroups, patient networks that are relatively similar (Jaccard index > 0.75) are mixed with patient networks with considerably larger differences (Jaccard index < 0.5). Moreover, this increasing diversity of patients is as expected most notable in larger random subgroups, evident from the negative correlation between subgroup size and average similarity (Pearson r = −0.58, p=0.019). In contrast, a positive correlation (Pearson r = 0.6, p=0.038) was instead observed for the Jaccard subgroups, demonstrating the effect of clustering using patient-specific topologies.

Altogether, we identified different subgroups with variation in subgroup size and heterogeneity that can inform distinct logic models. Our next step was to prepare the inputs for model optimization for each subgroup and investigate the effect of subgroup properties on model performance.

### PKN sizes after network reduction differ across subgroups

For each subgroup, we prepared combined PKNs (representing the assumed overlapping common network subset between patients) and corresponding training data. The PKNs were simplified with our two-threshold network reduction approach, with thresholds set manually for each subgroup to constraint sizes between 65 and 90 edges. We determined these sizes based on PKN sizes that still allowed for a feasible optimization. We also analyzed how cluster properties such as size (number of patients) and similarity (Jaccard index) are correlated and how these affect the PKN sizes with fixed thresholds across all subgroups. We defined two pairs of thresholds to represent a weak and a strong network reduction.

For the Jaccard subgroups, subgroup size and average similarity have a positive correlation with the combined PKN size (**Figure 4AB**). An interesting difference can be found between weak and strong reduction thresholds: subgroup similarity has a bigger effect on PKN size for the weak threshold (Pearson r=0.89, p=0.018) compared to the subgroup size (Pearson r=0.75, p=0.085). In contrast, subgroup size has a significantly higher positive correlation with PKN size for the strong threshold (Pearson r=0.84, p=0.037) compared to the subgroup similarity (Pearson r=0.36, p=0.049).

**Figure 4:**
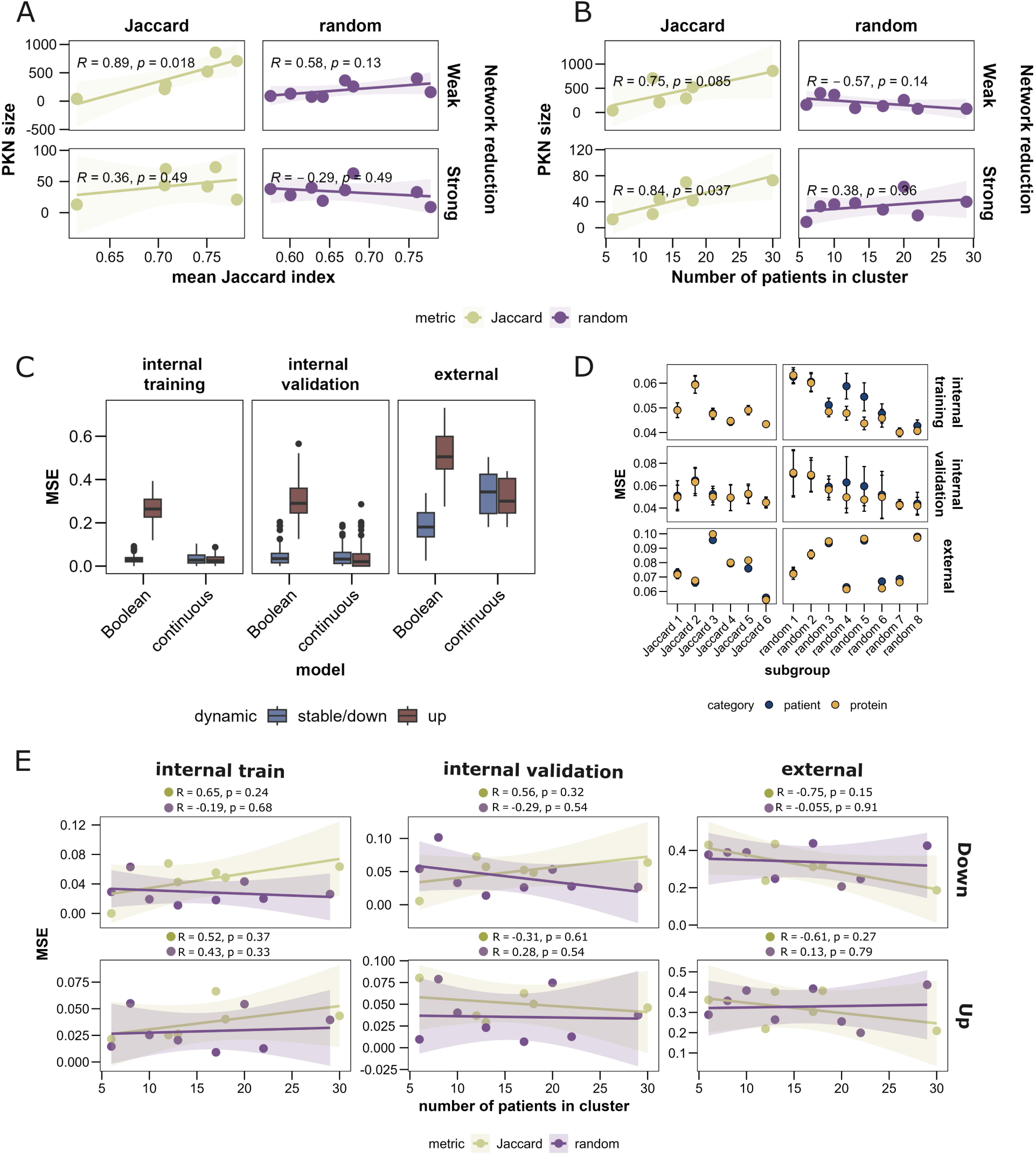
Optimizing models to different patient subgroups. **(A-B).** Analysis of the effect of network reduction on the Spearman correlation between PKN size and mean Jaccard index or number of patients in the cluster. A weak network reduction corresponds with threshold values of 2 and 1, while a strong network reduction with threshold values of 3 and 2 (see **Methods** for more detailed explanation on the thresholds). Every data point represents a model optimized to a different patient subgroup. A linear trendline is also included for each correlation analysis. **(C)**. Comparison of mean squared errors (MSE) across internal training (patients used for fitting), internal validation (patients within the same subgroup as training set, but excluded from training) and external set (patients from the cohort, not from same subgroup). Model simulations are divided into two dynamics: stable/down (Boolean/continuous) and up. Boxplots include all models for different subgroups and from a 5-fold crossvalidation with 10 repeats per fold. **(D).** Comparison MSE per patient with MSE per protein. The mean MSE across folds and repeats are given, with the bars indicating the standard deviation across optimizations. **(E)**. Analysis of the effect of subgroup size on the MSE. The mean of all optimizations for each subgroup is used to fit a linear trendline and calculate a Spearman correlation.

Generally, these trends suggest that with increasing subgroup size and similarity, a bigger number of overlapping proteins and edges can be identified across patients, resulting in a larger common PKN. This relation between subgroup size and PKN size is unique to the Jaccard subgroups, as with random subgroups and a weak reduction threshold, larger subgroups result in smaller combined PKN (Pearson r=-0.57, p=0.14). This observation confirms that in the larger random subgroups, less common mechanisms can be identified, which would be valuable for the dynamic model.

### Logic models capture protein activities across patients in a subgroup

For each subgroup, functional scores were converted to training data for both Boolean and logic-ODE models, with each patient represented as a distinct condition. We optimized the models using 5-fold cross-validation, and an ensemble of ten models was generated to account for stochastic variability. Model performance was evaluated on two datasets: the internal dataset (patients within the subgroup) and the external dataset (patients not included in the subgroup). Performance on the external dataset reflects the model’s ability to generalize to patients with less similarity to the subgroup. Comparing internal and external performance allows assessment of both patient specificity and subgroup representativeness. For our analyses, we classified protein activities into “dynamic categories” based on the final simulated values: up (1) or stable (0) for Boolean and up (> 0.55) or down (<0.45) for continuous.

We compared Boolean and continuous model performance for the Jaccard subgroups. For the Boolean models, mean squared error (MSE) was consistently higher for upregulated proteins than for stable proteins (**Figure 4C**). Training MSE was lowest, validation MSE was significantly higher across folds (p=2.83E-6 for up, p=0.019 for stable/down), and external MSE was highest. In contrast, continuous models showed similar MSE for both up- and downregulated dynamics, indicating less bias toward specific protein dynamics. Training and validation MSE also did not differ significantly (p=0.15 for up, p=0.34 for stable/down). Trends in training, validation, and external MSE were similar to those observed in Boolean models.

Our results highlight two complementary aspects of Boolean and logic-ODE models. Owing to their binary formulation, Boolean models cannot distinguish stable from downregulated states, as both are encoded as 0, and although upregulation can in principle be represented, Boolean optimisation tends to underestimate it by favouring zero states. In contrast, logic-ODE models operate in a continuous state space and explicitly capture graded up- and downregulation.

These advantages come at a computational cost: logic-ODE models required substantial simplification of the PKNs (maximum 100 edges) to allow feasible optimization and typically took hours to days to converge, whereas Boolean models could be optimized in minutes. Therefore, both modeling formalisms were essential within the COSMOS-CellNOpt integration, with Boolean models enabling efficient network refinement and logic-ODE models supporting detailed quantitative interpretation of patient-specific signalling dynamics. For this reason, all subsequent analyses are based on the logic-ODE models.

### Model performance per protein or per patient represent consistency of MSE across different conditions

To further study the model performance at the level of specific patients, proteins or sub-groups, we analyzed MSE per patient and per protein. Performance was generally consistent across patients and proteins (**Figure 4D**). The largest variation was observed in random subgroups, particularly in the validation set. For example, random subgroups 4–6 showed substantial variability in per-patient MSE, indicating that while the model fit some patients well, it struggled to capture others. This variability could be expected for random clusters, which include patients with divergent underlying mechanisms, previously shown with the variability in pairwise Jaccard indices (**Figure 3E**). These patients might not be adequately represented by the shared PKN and inputs, resulting in increased variation in MSE per patient. These findings highlight the advantage of Jaccard subgroups, where patients clustered by shared network mechanisms show more consistent model performance across individuals.

### Jaccard subgroup size have stronger correlations with model performance compared to random subgroups

We evaluated how patient subgroup size influences model performance by calculating Spearman correlation coefficients (R) between subgroup size (number of patients) and model performance (MSE) for the training, validation, and external datasets (**Figure 4E**). Although most correlations were not statistically significant, likely due to the small number of models per metric (6–8), the coefficients provide initial insight into potential relationships.

For the training dataset, correlations were generally positive, indicating that larger subgroup sizes tend to be associated with higher MSE (**Figure 4E**). This trend is consistent with the expectation that accurately fitting patients becomes more challenging as subgroup heterogeneity increases with bigger clusters. However, the magnitude of this effect was limited, as MSE values remained below 0.1 across subgroup sizes. Notably, for random subgroups, the correlation between subgroup size and MSE for downregulation was negative for both the training (r = −0.19, p = 0.68) and validation datasets (r = −0.29, p = 0.54), in contrast to the Jaccard subgroups.

The most pronounced differences between Jaccard and random subgroups were observed in the external dataset. Overall, MSE values for the external dataset (>0.1) were higher than those for the internal datasets (<0.1), which is expected given that the external patients were not included in the subgroup. Interestingly, Jaccard subgroups exhibited moderate to strong negative correlations between subgroup size and MSE (r = −0.75, p = 0.15 for downregulation and r = −0.61, p = 0.27 for upregulation; **Figure 4E**), whereas correlations for random subgroups were weak or negligible (r = −0.055, p = 0.91 and r = 0.13, p = 0.79, respectively).

While not statistically significant, this pattern suggests that increasing the size of subgroups defined through hierarchical clustering may capture shared biological characteristics that improve model generalization to patients outside the subgroup. Nevertheless, the consistently higher MSE observed for the external dataset emphasizes the importance of subgroup-specific modeling, as these models remain more representative of their target patient populations.

### Model error per patient and protein reveal areas of attention

We further analyzed optimized models to gain insights into regulatory mechanisms, focusing on the largest subgroup from hierarchical clustering. Model predictions were evaluated per protein and per patient by assessing whether up or down dynamics were correctly simulated. In general, different dynamics per patient were captured across proteins, with a few exceptions (**Figure 5A**). Standardized z-scores of prediction errors highlighted proteins with systematically higher errors. Training and validation z-scores were similar, with parent1 (regulator of DVL11, DVL21, DVL31, LRP51, and LRP61, see **Methods** for derivation of parent nodes) showing the highest z-scores (2.42 and 2.82 for train and validation, respectively; **Figure 5A**).

**Figure 5:**
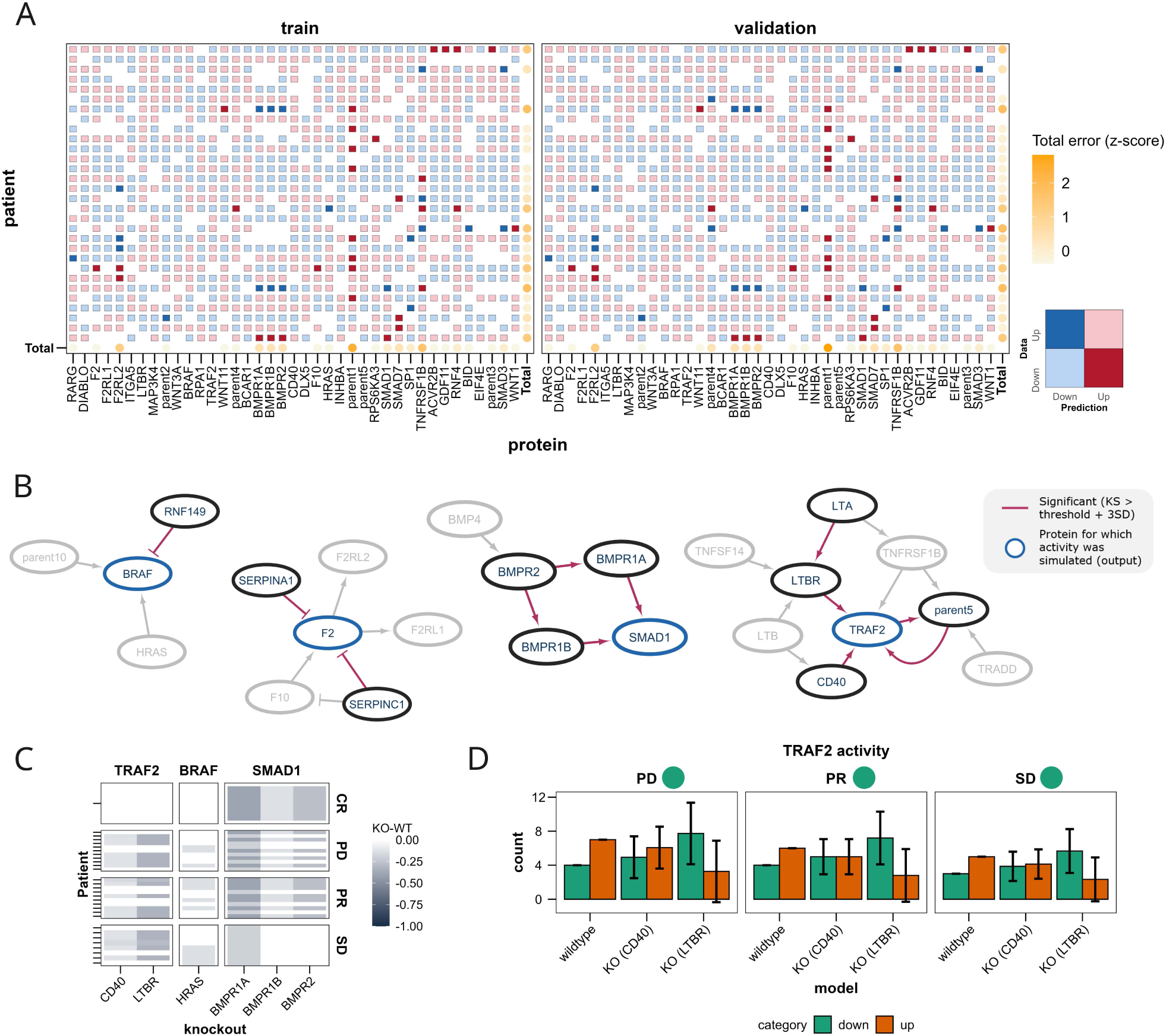
Biological analysis of an optimized model. **(A).** Error analysis for every protein (columns) and patient (rows), indicating for each protein and patient whether the up or down dynamic was predicted correctly. Additionally a standardized total error is given per patient or protein. White gaps represent missing data (i.e. patient had no functional score for this protein) or value from training was between 0.45 and 0.55 **(B).** Results from the sensitivity analyses, highlighting the sensitive interactions in the subnetworks of the PKN. **(C).** In silico knockout screening results, visualizing difference between knockout (KO) and wildtype (WT) model simulations for the proteins TRAF2, BRAF and SMAD1. The patients are categorized based on their responses (complete response, progressive disease, partial response and stable disease). A negative KO-WT indicates that the KO had a downregulating effect on the protein activity compared to the WT. **(D).** Influencing TRAF2 activity profiles for non-/partial responder patients through in silico knockouts. Every bar represents the mean count of for how many patients the protein is simulated as up or down across model optimization folds and repeats, and error bars indicate corresponding standard deviation. For each subplot, the color at the title represents the desired effect. For TRAF2, the aim was to increase the number of patients that exhibit a ‘down’ dynamic.

The node parent1 was consistently mispredicted as upregulated rather than downregulated for multiple patients. The activities of two upstream proteins of parent1 (WNT1, WNT3A) revealed how for the majority of both the train and validation sets, if both WNT1 and WNT3A were downregulated, parent1 was also correctly predicted as downregulation. For downregulated parent1 that were predicted as upregulation, a common occurrence was that WNT1 and / or WNT3A were upregulated, resulting in an incorrect upregulation of parent1. As the upstream nodes of WNT1 and WNT3A are already top nodes used to define patient conditions, the incorrect parent1 predictions could suggest that the top nodes for these specific patients might require some additional analyses to investigate whether there are incorrect or missing interactions related to these upstream proteins.

Additionally, BMPR1A, BMPR1B, and BMPR2 were mispredicted for three patients, with consistent error types across the three proteins. Since these proteins are functionally related, these errors also highlight a specific region of the network where the model predictions are less accurate and may require further refinement. Similar to the example of parent1, the network topology could be refined and analyzed for potential missing interactions.

### Parameter sensitivity analyses highlight relevant interactions in the protein network

We further analyzed the largest subgroup from hierarchical clustering by identifying interactions with the largest effect on protein activities. We focused on four proteins that are part of different, disconnected subnetworks with varying topologies to demonstrate how parameter sensitivity can identify relevant protein interactions: BRAF, F2, SMAD1 and TRAF2. We will also refer to these proteins as ‘outputs’ as we simulated these protein activities for our analyses. A multi-parametric sensitivity analysis (MPSA) was performed on the logic-ODE model. Through varying the interaction parameter k and analyzing its effects on the outputs, relevant protein interactions can be identified (see also **Methods**)[24].

Significant interactions were identified for each subnetwork corresponding to the outputs (**Figure 5B; Supplementary Figure 3**). Typically, direct interactions with the output protein were significant, though not all incoming interactions contributed equally. Identifying which interactions have the biggest impact on a protein of interest could be translated into potential targets for treatment or research. For example, BRAF has three incoming interactions, but only RNF149-BRAF was significant, while parent10-BRAF and HRAS-BRAF were not. This result creates a hypothesis that targeting RNF149-BRAF could affect BRAF more compared to parent10-BRAF or HRAS-BRAF. For SMAD1, both direct and upstream interactions were significant, including BMPR2-BMPR1A and BMPR2-BMPR1B. These results highlight the most influential interactions within the subgroup-specific model and identify key protein interactions for further analysis.

### Simulating knockouts allows for large-scale in silico screening

The model trained with this approach can be used to perform in silico knockouts. To mimic the knockout of a protein, we set all model parameters of its surrounding edges to 0, effectively excluding the protein from any network dynamics. As for the MPSA, we focused on the outputs BRAF, F2, SMAD1 and TRAF2. Most knockouts caused only minor changes in output activities. When knockout did affect the model, they generally led to decreased output activity (KO < WT), with changes ranging from 0.2 to 0.3 (**Figure 5C**).

To explore the functional relevance of these knockouts, we next focused on TRAF2 and whether in silico knockouts could shift protein activity profiles of partial or non-responders to resemble those of complete responders. Comparing complete responders (CR) with progressive disease (PD), partial responders (PR), and stable disease (SD) revealed that CRs had downregulated TRAF2 activity compared to a mix of up- and downregulation in PD, PR and SD. From the comprehensive in silico KO screening, CD40 and LTBR knockouts produced the largest effects on TRAF2, with several instances where TRAF2 activity after KO was lower than in the wildtype. Notably, LTBR knockout resulted in the strongest reductions as a larger number of patients were changed to a downregulated dynamic, bringing TRAF2 activity closer to the CR WT profile (**Figure 5D**). These findings suggest that both CD40 and LTBR could be targeted to reduce TRAF2 activity, with LTBR expected to have the stronger effect. Altogether, these results demonstrate how in silico knockouts enable more detailed analyses of optimized models. Beyond reconstructing regulatory mechanisms, they can guide the selection of targets to achieve specific changes in protein activity within a patient subgroup.

### TCGA-KIRC clusters show distinct characteristics in both biological function enrichment and immune subtypes

To evaluate the scalability of our method, we next applied the workflow to a larger cohort. Transcriptomic data were obtained from the TCGA-KIRC project [25]. After restricting the dataset to patients with primary tumor samples, 533 cases remained. MOON was unable to generate patient-specific networks for seven patients, resulting in a final cohort of 526 patients. We clustered these patients into six subgroups based on pairwise Jaccard indices, yielding cluster sizes of 239, 145, 86, 36, 14 and 6 patients for clusters 1 - 6, respectively. We then performed enrichment analysis as described for the previous dataset (**Figure 3**). Due to its small size (N = 6), cluster 6 was excluded from downstream enrichment analyses. The resulting enrichment scores revealed both shared and distinct biological features among clusters (**Figure 6A**). For example, cluster 4 showed a lower enrichment score (score = 0.73) for epithelial mesenchymal transition compared to the other clusters (score > 4.5), a pathway commonly associated with metastatic potential.

**Figure 6:**
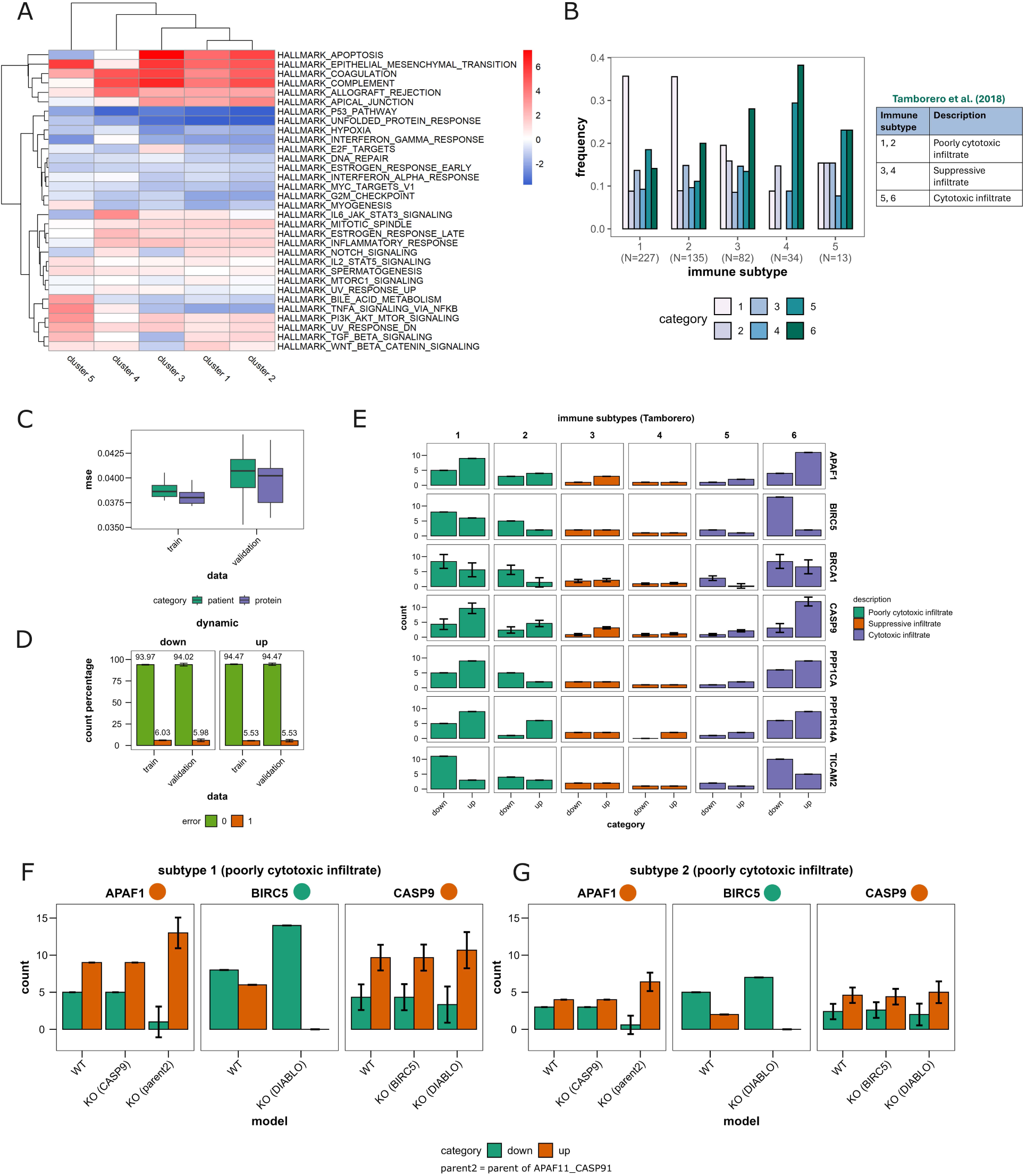
TCGA KIRC application. **(A).** Enrichment analysis of subgroups created from the TCGA KIRC cohort. Enrichment scores are provided for different HALLMARK gene sets for five different clusters. Cluster 6 was excluded for this visualization due to the small cluster size (N=6). **(B).** Immune subtype frequencies per cluster. The immune subtypes from Tamborero et al. (2018) [27] are visualized with frequencies scaled by cluster size. For each cluster, the cluster size (N) is also provided. Moreover, a small table with immune subtype definitions is added. **(C).** Model fitting results for the training and validation patients. Boxplots represent optimizations across different folds and repeats. **(D).** The accuracy of model predictions for different dynamic categories (up/down). For each correct (error=0) and incorrect (error=1) prediction, a percentage is visualized to represent what fraction of the total these correct/incorrect predictions cover. Error bars represent deviation across different optimizations. **(E).** Frequencies of up/down dynamics across different immune subtypes and selected proteins. The detailed description of each immune subtype is indicated by color. The mean counts are given, with the error bars visualizing the standard deviation across repeats. **(F-G).** Analysis of in silico knockouts and their effects on protein activities for immune subtypes 1 and 2. The indicated color at each subplot title indicates the dynamics category that is preferred based on the favorable immune subtype 6. For example, the aim of APAF1 is to increase the count of “up” dynamics through the knockouts. .

We next examined whether the identified subgroups corresponded to established patient annotations or phenotype-based classifications. Since the enrichment analysis revealed distinct biological features among clusters, we hypothesized that these differences might also be reflected in phenotypic signatures. Several previous studies have used transcriptomic profiles to define immune-related tumor subtypes, either based on broad tumor microenvironmental and survival characteristics [26] or on more specific features such as immune cell infiltration [27] and predicted response to immunotherapy [28]. Using the subtype annotations from these three studies (Thorsson, Tamborero, Bagaev), we assessed their distributions across our clusters (**Figure 6B, Supplementary Figure 4A**). Overall, most subgroups were enriched for the Thorsson C3 subtype, indicative of an inflammatory phenotype. However, more pronounced differences among subgroups emerged when comparing the Bagaev and Tamborero classifications, suggesting that these subtypes may capture finer-grained immunological variation across clusters.

Notably, cluster 4 exhibited higher frequencies of Tamborero subtypes 5 and 6, both classified as cytotoxic infiltrate, which may reflect a more favorable immune landscape within these tumors. This observation is consistent with the positive enrichment score for the interferon_gamma_response pathway in cluster 4 (score = 0.22; **Figure 6A**), compared to negative scores in the other clusters (score < −1.85), highlighting a potential role for immune infiltration. Overall, although these subgroups were defined through a data-driven, network-based approach using COSMOS on omics data, the results suggest that they capture meaningful clinically relevant distinctions in this dataset.

### Optimized logic-ODE models show good agreement in capturing up and down dynamics

Next, we focused on cluster 4, due to its distinct characteristics identified in the cluster analyses (**Figure 6A-B**). Aggregating the patient-specific MOON networks produced a PKN of 108 nodes and 89 edges. Consistent with our approach for the SU2C-MARK cohort, we optimized models using 5-fold crossvalidation. To account for stochasticity, a total of 15 models were generated.

The optimized models had similar MSE for training and validation across folds (**Figure 6C**). Error distributions across patients and proteins were also consistent, indicating a robust overall fit. We also evaluated the model’s accuracy in predicting the direction of activities (“up” (activity > 0.5) or “down” (activity < 0.5)). High prediction accuracy (> 93%) was observed across both dynamics and training/validation data (**Figure 6D**). These results suggest that, while the exact protein activity values may not be perfectly simulated, the model successfully captures the general dynamic behavior of the subgroup and differentiates patient-specific responses.

### In silico knockouts highlight proteins that adjust protein activities to be more similar to cytotoxic infiltrate immune subtypes

Next, we conducted an in silico knockout screening of all proteins in the network that were not top nodes used to define individual patients. The goal was to identify proteins whose perturbation could shift a patient profile with less favorable prognostic or therapeutic dynamics toward a more responsive state. We focused on the Tamborero immune subtypes, given that cluster 4 exhibited increased frequencies of subtypes 5 and 6 (cytotoxic infiltrate), which are generally associated with improved response to immunotherapy. Using the complete in silico knockout screening, we identified proteins for which knockouts induced a substantial change in activity (> 0.1; **Supplementary Figure 4B**). We further refined this list by focusing on proteins that vary the most across patients. We considered the standard deviation of protein activities across patients, selecting only proteins with a standard deviation of at least 0.25 (**Supplementary Figure 4C**). This combined approach yielded seven candidate proteins: APAF1, BIRC5, BRCA1, CASP9, and PPP1CA, PPP1R14A and TICAM2.

We first examined the activities of the selected proteins across immune subtypes in greater detail, categorized as “up” or “down” (**Figure 6E**). For example, in immune subtype 6, APAF1 and CASP9 were predominantly upregulated, whereas BIRC5 and TICAM2 were more frequently downregulated. In contrast, the poorly cytotoxic infiltrate subtypes (1 and 2) displayed different patterns: BIRC5 and CASP9 activities are more evenly split between up and down. Based on these observations, we then screened the in silico knockouts to identify targets in the poorly cytotoxic infiltrate subtypes that could shift APAF1 and CASP9 toward upregulation while promoting downregulation of BIRC5.

The in silico screening identified promising targets to shift protein activities in subtype 1 toward patterns observed in subtype 6 (**Figure 6F**). For APAF1, only the knockout of parent_of_APAF11_CASP91 (parent2) substantially altered APAF1 dynamics, changing the counts of up- and downregulated states. Notably, parent2 knockout resulted in nearly all patients exhibiting upregulated APAF1, more closely resembling the majority of subtype 6 patients. Similarly, DIABLO knockout induced changes in the dynamics of BIRC5 and CASP9.. These results demonstrate that in silico knockouts can help identify proteins of interest, such as DIABLO, for further investigation. These knockouts had similar effects for patients with subtype 2, another immune subtype characterized as poorly cytotoxic infiltrate (**Figure 6G**). These findings highlight the benefits of incorporating predefined immune subtypes for post-hoc analyses such as in silico knockout screenings, providing a framework for hypothesis generation.

## Discussion

We developed a workflow that derives dynamic logic models directly from static omics data. Using COSMOS, the method first constructs static, patient-specific protein networks from which corresponding functional scores are derived[8]. Next, the workflow generates CellNOpt-compatible inputs, including patient clustering and network simplification. This addition enabled successful fitting of dynamic Boolean and logic-ODE models to transcriptomic cohorts such as SU2C-MARK lung cancer and TCGA-KIRC[22,25]. Even though these data sets include only one measurement per patient, leveraging inter-patient variability allowed us to obtain consensus mechanistic models that remain capable of simulating patient-specific responses. In other words, our workflow expands the applicability of the logic models in CellNOpt to patient data that do not include multiple perturbations.

A major bottleneck in applying dynamic logic models at scale is the time-consuming manual curation of prior knowledge networks. To support scalability and adaptability, we focused on automating this step by leveraging the abundance of available high-throughput omics data. In contrast to previous studies that integrate omics data by mapping them onto pre-existing, manually curated models [11,14], our approach infers signaling networks directly from omics data. Integrating these data-driven prior knowledge networks with CellNOpt enabled the inference of dynamic behavior of underlying regulatory systems.

A key advantage of the resulting dynamic models is their interpretability and ability to perform in silico perturbations, which are valuable for advancing precision oncology[3]. We demonstrated that simulated knockouts can shift samples from poor to favorable phenotypic or immune profiles, supporting the identification and prioritization of therapeutic targets. Additionally, the application to the TCGA-KIRC dataset demonstrates the applicability of our workflow to a larger cohort. These results provide opportunities to explore consensus among patients within a subgroup while capturing patient-specific dynamics, offering a framework for further mechanistic studies and guiding hypothesis-driven experimental design.

Additionally, central to our approach is the clustering of patients based on MOON-derived network topology, without requiring clinical or molecular annotations. These topology-based subgroups showed biologically meaningful distinctions through enrichment analyses and immune subtype associations, and produced more coherent combined networks than random subgroups. While topology alone already captures important variation, integrating additional patient-specific information can further enhance modeling. For example, subgroups defined by tumor grade, pathologic stage, or predefined immune subtypes reflect differences in prognosis and treatment response. Investigating the dynamics of each category with separate dynamic models could uncover distinct regulatory mechanisms. Similarly, in silico perturbation analyses can reveal mechanistic differences across stages or subtypes, potentially highlighting strategies to restore aberrant signaling toward healthy dynamics[29].

Patient-specific driver mutations offer another complementary layer of information. Although cancer cells accumulate numerous genetic alterations that can shape individual responses to therapy, only a subset of mutations critically influence tumor progression[30]. Unfortunately, many driver mutations are unknown. As experimental identification can be challenging due to the sheer size of the search space, in silico analyses can help identify candidate driver proteins by detecting deviating signaling dynamics within or across subgroups [30].

While our workflow can be readily applied to diverse datasets and case studies, several components can be expanded or adapted. Although we did not further employ the Boolean model due to limitations on its simulated dynamics, simpler approaches may be advantageous in certain scenarios. Boolean models require substantially less computational power and may suffice when protein activities do not need to be fully continuous. If Boolean models are too simplistic and continuous models too complex, an intermediate option could be considered such as multi-state or fuzzy logic models. These approaches allow proteins to adopt more than two states, which is more representative of signaling dynamics[19,31].

Future implementations within the workflow could also incorporate alternative subgroup network inference tools, such as CORNETO, to infer combined networks directly from omics data[32]. Such approaches may help alleviate computational bottlenecks of our current clustering approach like pairwise Jaccard index calculations. Therefore, an important next step would be to enable benchmarking of different network curation heuristics through for instance integration into platforms such as NetworkCommons [33]. Furthermore, applying the workflow to large-scale datasets, including single-cell atlases, could be promising avenues for studying shared and distinct mechanisms in patients[34] These datasets enable analysis of heterogeneity across cell populations, and as single-cell technologies advance, they provide opportunities to model cell-type-specific dynamics in silico[35,36]. Addressing computational and modeling challenges in this context will be crucial for extending the scalability and applicability of our method.

Altogether, the flexibility of our approach to integrate diverse datasets and modeling approaches highlights the modular design we envision. By linking omics-derived networks through tools such as COSMOS and CellNOpt, we provide a solution to two challenges in dynamic logic modeling: the automated definition of model structure and the integration of high-throughput omics data for model building and optimization. This framework supports patient stratification and the investigation of patient-specific signaling dynamics, providing insights that can guide the identification of anti-cancer drug targets and the design of personalized treatment strategies. As computational and experimental tools continue to advance rapidly, modular, data-driven approaches that combine multiple tools will be increasingly valuable for precision oncology.

## Materials and methods

### Prostate cancer TCGA data

We tested the automated protein network curation using 483 human prostate cancer patients from the TCGA data set [18], the same data set used by Montagud et al. (2022) [11]. We retrieved the RNAseq data (RSEM, Batch normalized from Illumina HiSeq_RNASeqV2) from https://www.cbioportal.org/study/summary?id=prad_tcga_pan_can_atlas_2018.

### Lung cancer cohort data

We used the SU2C-MARK lung cancer patient cohort for testing of the workflow. This data set contains patients with advanced non-small cell lung cancer treated with an anti-PD-1/PD-L1 agent. Measurements were derived from profiling tumor tissue with whole transcriptome RNA-seq [22]. We performed log2 transformation and normalized the RNA-seq data across patients. Next, we utilized CollecTRI to estimate transcription factor (TF) activities [37]. Additionally, patients were annotated with the name of the agent, tumor histology and treatment response (best overall response; BOR). There were significant differences between TF activities of several subsets based on these annotations. We therefore decreased the variability across patients by focusing on the patients treated with the anti-PD-1 agent nivolumab. This subset consisted of 60 patients with adenocarcinoma (71%), squamous cell carcinoma (23%), large cell neuroendocrine carcinoma (1%) and other histologies (4%). Three types of BOR were recorded: complete / partial response (30%), stable disease (29%) and progressive disease (41%) [22].

### TCGA-KIRC data

The second data set we acquired is the transcriptomics data from the Cancer Genome Atlas Kidney Renal Clear Cell Carcinoma (TCGA-KIRC) data collection [25]. The data was filtered to only contain patients where the recorded data was from a primary tumor, resulting in a data set consisting of 526 patients. Moreover, immune subtypes for the filtered TCGA-KIRC patients were derived from multiple resources. Firstly, we utilized the immune subtypes defined by Bagaev et al. (2021), immune-enriched fibrotic, immune-enriched non-fibrotic, fibrotic and desert [28]. These categories are classified based on transcriptomic analyses and correlate with patient response to immunotherapy. Secondly, immune subtypes focusing on immune infiltration defined by Tamborero et al. (2018) were included. They used the relative abundances of immune cell populations to distinguish immune infiltration patterns and categorize patients into six different phenotypes [27]. Lastly, the study by Thorsson et al. (2018) identified six immune subtypes (C1 - C6) based on different factors such as macrophage:lymphocyte ratio, Th1:Th2 ratio, proliferation rates and intratumoral heterogeneity [26]. As each of these immune subtypes were defined based on different metrics, we included all three for analysis of subtype frequencies per subgroup.

### Deriving contextualized mechanistic models

COSMOS requires both a prior knowledge network (PKN) and TF activities. Within the PKN, each node represents a protein and each edge a causal interaction between linked proteins. We derived the PKN from Omnipath [38]. The generic PKN was further processed in three steps: **(1)** Filtering of the network to keep only TF proteins that are significantly expressed, which we defined as an absolute activity larger than 1.5. **(2)** Reduction through removing proteins that are not connected to any TF. **(3)** Correction to remove any edge that leads to an incoherence between TF and its target. Additionally, MOON, which is part of COSMOS, uses an univariate linear model (ULM) to estimate activities of upstream proteins based on TF activities. MOON proceeds to extract a patient-specific network from the PKN using the functional scores.

Thresholds on protein activity values and removing parts of the network with inconsistencies across functional scores reduce the PKN into a contextualized, patient-specific MOON network [8]. This simplification of the PKN also includes merging nodes (parents) that share the same downstream proteins (children). These nodes are compressed into general ‘parent’ nodes.

We used the Jaccard index to define similarity between patient-specific MOON networks, before clustering the networks through hierarchical clustering with complete linkage. The Jaccard index is defined as 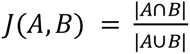 with A and B being the set of edges of two different patients. Patient-specific contextualized networks were aggregated (i.e. union of all networks) across patients clustered into the same subgroup. As resulting combined protein networks can have several thousands of edges, we defined a two-threshold reduction approach to simplify these networks.

Firstly, a selection of proteins is created based on a predefined first threshold. This selection is used to create a subnetwork from the original MOON network, so the all edges satisfy either of the two requirements: (1) both source and target are selected proteins based on the first threshold, or (2) either source or target is part of selected proteins while the other protein is an intermediate between two selected proteins. Next, a second threshold (less strict than first threshold) is used to further refine edges selected based on requirement (2): all nodes within the simplified network are required to have an absolute functional score above the second threshold. This method simplifies networks significantly while taking into account protein neighbors to preserve as large as possible subnetworks as possible.

Lastly, we used node and edge filtering to address issues that may arise from combining patient-specific networks and data into one model, including data sparsity and inconsistencies. Nodes lacking data in ≥20% of subgroup patients were excluded, reducing sparsity in the training data. Additionally, patient-specific networks and corresponding functional scores have been filtered to remove incoherent signs between proteins. However, combining networks risks introducing new incoherent interactions in the network, which complicate model fitting. We therefore removed edges which are not present in ≥30% of patient-specific networks. The partial removal still ensures that there are patient-specific interactions present, while improving model optimization.

### Enrichment analysis

Gene sets such as the hallmark collection of MSigDB [23], link biological functions to genes. We however clustered based on a Jaccard index determined with network edges. Therefore, we determined biological functions per edge. To assign a biological function to an edge, we set a requirement that for an edge of protein A and B, both proteins are assigned to this function in the hallmark gene set. For the enrichment analysis, we counted the number of times an edge appeared in each cluster across all patients and normalized by cluster size. We performed enrichment analysis separately for the Jaccard and random subgroups using the ULM approach of decoupleR [39]. In other words, resulting enrichment scores are for instance relative to other Jaccard subgroups, but not random subgroups.

### Logic modeling with CellNOpt

For our logic modeling, we used CellNOpt, an open-source R software package featuring different logic formalisms. One method of distinguishing the different formalisms is based on model state, ranging from binary (Boolean) to continuous (logic-ODE). The different formalisms adhere to a similar pipeline consisting of **(1)** Importing data, **(2)** Network processing, and **(3)** Model training [19].

For the binary model, we utilized a Boolean (ON/OFF) model which is optimized with the built-in genetic algorithm of CellNOpt [19]. We encoded functional scores into different categories (upregulated / stable) using two values (initial and final) to create a two-state binary model. For the continuous model, we created logic-ODE models with the add-on CNORode of CellNOpt [12]. This logic-ODE model is based on the transformation described by Wittman et al. (2009) going from Boolean to continuous models [40]. Model optimization is performed with MEIGOR and its implemented metaheuristic enhanced scatter search [41]. For our analyses and distinction between dynamics, we defined upregulation as protein activities >0.6 and downregulation as activities <0.45.

Both the Boolean and continuous model require two inputs: a prior knowledge network (PKN) and experimental data. These are given through a SIF and MIDAS file, respectively. The patient-specific MOON networks derived with COSMOS were combined into one PKN for CellNOpt. We utilized the functional scores inferred with MOON for the MIDAS file. The MIDAS format makes a distinction between whether a value represents a treatment (TR), data acquisition (DA) or data value (DV). With TR, we indicate for which patient training data are given and which proteins are active as input nodes. Binary values (0 = not present, 1 = present) are used to indicate in which conditions patient and corresponding inputs are present. We used DA values to distinguish between the two simulation states, indicating whether protein activities are initial values (t=0) or final values (t=10). Lastly, DV contains processed training data values for each patient and protein. Depending on the model state, we either converted the activities into binary values (ON = 1, OFF = 0) or scaled values to a range between 0 and 1.

### Logic-ODE parameter analyses

Three parameters can be optimized in the logic-ODE formalism: τ_i_ (life-time of node i), n_ij_ (Hill coefficient of interaction between nodes i and j) and k_ij_ (strength of regulation of node j on i) [19]. We fixed n_ij_ at 3, and focused post-hoc analyses on the k_ij_ parameter. Firstly, we performed multi-parametric sensitivity analysis (MPSA) [24]. This global sensitivity analysis allows us to investigate for specified proteins which protein interactions have the largest impact on the simulated protein activity by varying the k parameter through Latin hypercube sampling.

Differences in output activities are analyzed with the Kolmogorov-Smirnov (KS) statistic, where higher values represent a higher impact of the parameter on the model output. We performed three independent runs with 20,000 parameter samples (maximum ±0.3 variation. Next, we performed in silico knockout screenings. We mimicked protein knockout by setting all kij parameters of interactions linked to the knockout protein to 0. Results of this adjusted model (KO) can be compared with the original model (WT) for investigating the effect of knockouts on simulated protein activities.

### Statistical analyses

If not mentioned otherwise, we use a significance threshold of 5% for statistical tests. Several types of statistical tests were utilized, such as ANOVA for differences between groups, Spearman for correlation and pairwise Fisher’s exact test with Benjamini-Hochberg correction for differences in proportions of categories across two groups.

Specifically for the analyzing model optimization, model fit was evaluated through calculating the mean squared error (MSE) between training and simulated data. To look at possible correlations between subgroup properties and model fit, we utilized the Spearman’s rank correlation coefficient to analyze the monotonic relationship between the two variables. Additionally, we fitted linear trend lines and determined 95% confidence intervals of the regression line.

## Acknowledgements

This work was supported by the Netherlands Organization for Scientific Research (NWO) Gravitation program IMAGINE! (project number 24.005.009) and by NWO Aspasia grant (project number 015.021.065) awarded to F.E., which supported B.W.. We acknowledge funding to J.S.R. by the German Federal Ministry of Education and Research (Bundesministerium für Bildung und Forschung BMBF) to support A.D.. Y.B. acknowledges the support of the Heidelberg Faculty of Medicine at Heidelberg University through the Medical Data Scientist Fellowship. The results published here are in whole or in part based upon data generated by The Cancer Genome Atlas (TCGA) Research Network. ChatGPT was used to assist in language editing and improving readability. The authors reviewed and verified all generated content.

## Author contributions

B.W.: Formal analysis, Investigation, Methodology, Software, Validation, Visualization, Writing – original draft, Writing - review & editing

Y.B.: Software, Validation

J.S.R.: Conceptualization, Funding acquisition, Writing – review & editing

F.E.: Funding acquisition, Project administration, Supervision, Writing – review & editing

A.D.: Conceptualization, Methodology, Project administration, Supervision, Writing – review & editing

## Code availability

All code to ensure reproducibility of the results is available at https://github.com/saezlab/ficus. This includes vignettes to data preprocessing, preparation of COSMOS and CellNOptR inputs, optimizing logic models and running the cluster enrichment analyses and knockout simulations. The repository can be installed as the R package FICUS.

## References

1. Proietto M, Crippa M, Damiani C, Pasquale V, Sacco E, Vanoni M, et al. Tumor heterogeneity: preclinical models, emerging technologies, and future applications. Front Oncol. 2023;13:1164535.

2. Liaki V, Barrambana S, Kostopoulou M, Lechuga CG, Zamorano-Dominguez E, Acosta D, et al. A targeted combination therapy achieves effective pancreatic cancer regression and prevents tumor resistance. Proc Natl Acad Sci U S A. 2025;122:e2523039122.

3. Marques L, Costa B, Pereira M, Silva A, Santos J, Saldanha L, et al. Advancing precision medicine: A review of innovative in silico approaches for drug development, clinical pharmacology and personalized healthcare. Pharmaceutics. 2024;16:332.

4. Mangul S, Martin LS, Hill BL, Lam AK-M, Distler MG, Zelikovsky A, et al. Systematic benchmarking of omics computational tools. Nat Commun. 2019;10:1393.

5. Dugourd A, Saez-Rodriguez J. Footprint-based functional analysis of multiomic data. Curr Opin Syst Biol. 2019;15:82–90.

6. Dai X, Shen L. Advances and trends in omics technology development. Front Med (Lausanne). 2022;9:911861.

7. Cancer Genome Atlas Research Network. Comprehensive molecular characterization of clear cell renal cell carcinoma. Nature. 2013;499:43–9.

8. Dugourd A, Kuppe C, Sciacovelli M, Gjerga E, Gabor A, Emdal KB, et al. Causal integration of multi-omics data with prior knowledge to generate mechanistic hypotheses. Mol Syst Biol. 2021;17:e9730.

9. Dugourd A, Lafrenz P, Mañanes D, Paton V, Fallegger R, Kroger A-C, et al. Modeling causal signal propagation in multi-omic factor space with COSMOS [Internet]. bioRxiv. 2024. Available from: 10.1101/2024.07.15.603538

10. Garrido-Rodriguez M, Zirngibl K, Ivanova O, Lobentanzer S, Saez-Rodriguez J. Integrating knowledge and omics to decipher mechanisms via large-scale models of signaling networks. Mol Syst Biol. 2022;18:e11036.

11. Montagud A, Béal J, Tobalina L, Traynard P, Subramanian V, Szalai B, et al. Patient-specific Boolean models of signalling networks guide personalised treatments. Elife [Internet]. 2022;11. Available from: 10.7554/eLife.72626

12. Eduati F, Jaaks P, Wappler J, Cramer T, Garnett MJ, Saez-rodriguez J, et al. Patient-specific logic models of signaling pathways from screenings on cancer biopsies to prioritize personalized combination therapies. 2020;1–13.

13. Fröhlich F, Gerosa L, Muhlich J, Sorger PK. Mechanistic model of MAPK signaling reveals how allostery and rewiring contribute to drug resistance. Mol Syst Biol. 2023;19:e10988.

14. Béal J, Montagud A, Traynard P, Barillot E, Calzone L. Personalization of logical models with multi-omics data allows clinical stratification of patients. Front Physiol. 2018;9:1965.

15. Gjerga E, Trairatphisan P, Gabor A, Koch H, Chevalier C, Ceccarelli F, et al. Converting networks to predictive logic models from perturbation signalling data with CellNOpt. Bioinformatics. 2020;36:4523–4.

16. Ruscone M, Tsirvouli E, Checcoli A, Turei D, Barillot E, Saez-Rodriguez J, et al. NeKo: A tool for automatic network construction from prior knowledge. PLoS Comput Biol. 2025;21:e1013300.

17. Heo YJ, Hwa C, Lee G-H, Park J-M, An J-Y. Integrative multi-omics approaches in cancer research: From biological networks to clinical subtypes. Mol Cells. 2021;44:433–43.

18. Hoadley KA, Yau C, Hinoue T, Wolf DM, Lazar AJ, Drill E, et al. Cell-of-origin patterns dominate the molecular classification of 10,000 tumors from 33 types of cancer. Cell. 2018;173:291–304.e6.

19. Terfve C, Cokelaer T, Henriques D, MacNamara A, Goncalves E, Morris MK, et al. CellNOptR: A flexible toolkit to train protein signaling networks to data using multiple logic formalisms. BMC Syst Biol [Internet]. 2012;6. Available from: 10.1186/1752-0509-6-133

20. Türei D, Valdeolivas A, Gul L, Palacio-Escat N, Klein M, Ivanova O, et al. Integrated intra-and intercellular signaling knowledge for multicellular omics analysis. Mol Syst Biol. 2021;17:e9923.

21. Schubert M, Klinger B, Klünemann M, Sieber A, Uhlitz F, Sauer S, et al. Perturbation-response genes reveal signaling footprints in cancer gene expression. Nat Commun. 2018;9:20.

22. Ravi A, Hellmann MD, Arniella MB, Holton M, Freeman SS, Naranbhai V, et al. Genomic and transcriptomic analysis of checkpoint blockade response in advanced non-small cell lung cancer. Nat Genet. 2023;55:807–19.

23. Liberzon A, Birger C, Thorvaldsdóttir H, Ghandi M, Mesirov JP, Tamayo P. The Molecular Signatures Database (MSigDB) hallmark gene set collection. Cell Syst. 2015;1:417–25.

24. Zi Z. Sensitivity analysis approaches applied to systems biology models. IET Syst Biol. 2011;5:336–46.

25. Akin O, Elnajjar P, Heller M, Jarosz R, Erickson BJ, Kirk S, et al. The Cancer Genome Atlas Kidney Renal Clear Cell Carcinoma collection (TCGA-KIRC) [Internet]. The Cancer Imaging Archive; 2016. Available from: 10.7937/K9/TCIA.2016.V6PBVTDR

26. Thorsson V, Gibbs DL, Brown SD, Wolf D, Bortone DS, Ou Yang T-H, et al. The immune landscape of cancer. Immunity. 2018;48:812–30.e14.

27. Tamborero D, Rubio-Perez C, Muiños F, Sabarinathan R, Piulats JM, Muntasell A, et al. A pan-cancer landscape of interactions between solid tumors and infiltrating immune cell populations. Clin Cancer Res. 2018;24:3717–28.

28. Bagaev A, Kotlov N, Nomie K, Svekolkin V, Gafurov A, Isaeva O, et al. Conserved pan-cancer microenvironment subtypes predict response to immunotherapy. Cancer Cell. 2021;39:845–65.e7.

29. Cheng YQ, Wang SB, Liu JH, Jin L, Liu Y, Li CY, et al. Modifying the tumour microenvironment and reverting tumour cells: New strategies for treating malignant tumours. Cell Prolif. 2020;53:e12865.

30. Ostroverkhova D, Przytycka TM, Panchenko AR. Cancer driver mutations: predictions and reality. Trends Mol Med. 2023;29:554–66.

31. Wynn ML, Consul N, Merajver SD, Schnell S. Logic-based models in systems biology: a predictive and parameter-free network analysis method. Integr Biol (Camb). 2012;4:1323–37.

32. Rodriguez-Mier P, Garrido-Rodriguez M, Gabor A, Saez-Rodriguez J. Unifying multi-sample network inference from prior knowledge and omics data with CORNETO. Nat Mach Intell. 2025;7:1168–86.

33. Paton V, Türei D, Ivanova O, Müller-Dott S, Rodriguez-Mier P, Venafra V, et al. NetworkCommons: bridging data, knowledge, and methods to build and evaluate context-specific biological networks. Bioinformatics [Internet]. 2025;41. Available from: 10.1093/bioinformatics/btaf048

34. Pan L, Parini P, Tremmel R, Loscalzo J, Lauschke VM, Maron BA, et al. Single Cell Atlas: a single-cell multi-omics human cell encyclopedia. Genome Biol. 2024;25:104.

35. Guo W, Chen Z, Gu J. Computational modeling of single-cell dynamics data. Brief Bioinform [Internet]. 2025;26. Available from: 10.1093/bib/bbaf305

36. Guo X, Adelaja A, Singh A, Wollman R, Hoffmann A. Modeling heterogeneous signaling dynamics of macrophages reveals principles of information transmission in stimulus responses. Nat Commun [Internet]. 2025;16. Available from: 10.1038/s41467-025-60901-3

37. Müller-Dott S, Tsirvouli E, Vazquez M, Ramirez Flores RO, Badia-I-Mompel P, Fallegger R, et al. Expanding the coverage of regulons from high-confidence prior knowledge for accurate estimation of transcription factor activities. Nucleic Acids Res. 2023;51:10934–49.

38. Türei D, Korcsmáros T, Saez-Rodriguez J. OmniPath: guidelines and gateway for literature-curated signaling pathway resources. Nat Methods. 2016;13:966–7.

39. Badia-I-Mompel P, Vélez Santiago J, Braunger J, Geiss C, Dimitrov D, Müller-Dott S, et al. decoupleR: ensemble of computational methods to infer biological activities from omics data. Bioinform Adv. 2022;2:vbac016.

40. Wittmann DM, Krumsiek J, Saez-Rodriguez J, Lauffenburger DA, Klamt S, Theis FJ. Transforming Boolean models to continuous models: Methodology and application to T-cell receptor signaling. BMC Syst Biol. 2009;3:98–98.

41. Egea JA, Henriques D, Cokelaer T, Villaverde AF, MacNamara A, Danciu DP, et al. MEIGO: An open-source software suite based on metaheuristics for global optimization in systems biology and bioinformatics. BMC Bioinformatics. 2014;15:1–9.

