## Supplementary Information for "Combining prior knowledge and transcriptomics data for logic models of patient subgroups"

**Supplementary data**

Authors

Bi-rong Wang^abc^

Yunfan Bai^c^

Julio Saez-Rodriguez*^cd^

Federica Eduati*^ab^

Aurélien Dugourd*^cd^

*=corresponding author

Author affiliations

^a^Department of Biomedical Engineering, Eindhoven University of Technology, PO Box 513, Eindhoven 5600MB, the Netherlands

^b^Institute for Complex Molecular Systems, Eindhoven University of Technology, 5600 MB Eindhoven, The Netherlands

^c^Institute for Computational Biomedicine, Heidelberg University Hospital, Heidelberg, Germany

^d^European Bioinformatics Institute, EMBL, Hinxton, UK

Contact information

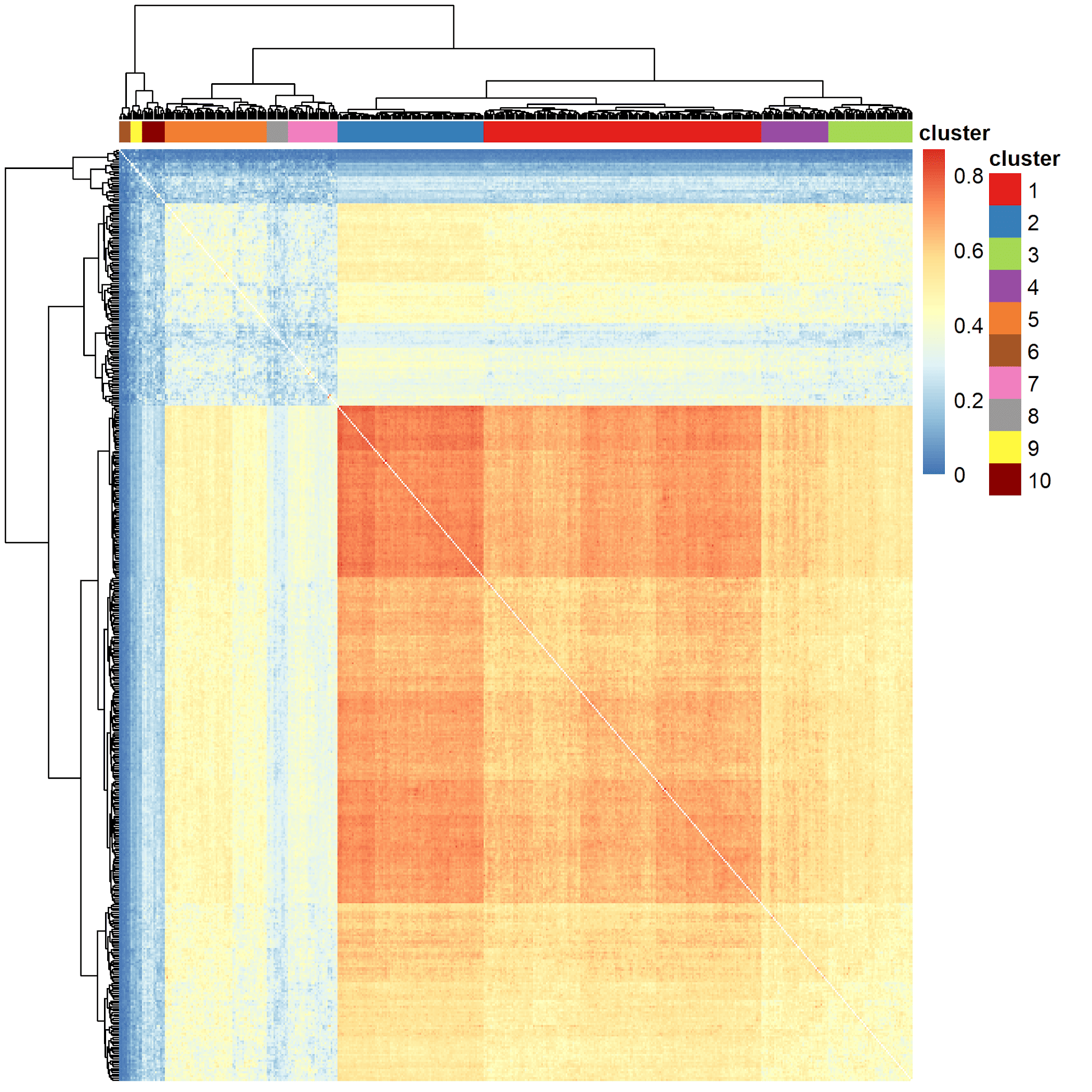

**Supplementary Figure 1 :** Heatmap of pairwise Jaccard indices across patients of the TCGA prostate cancer data set (x and y axes represent patients). Corresponding cluster index per patient is also indicated.

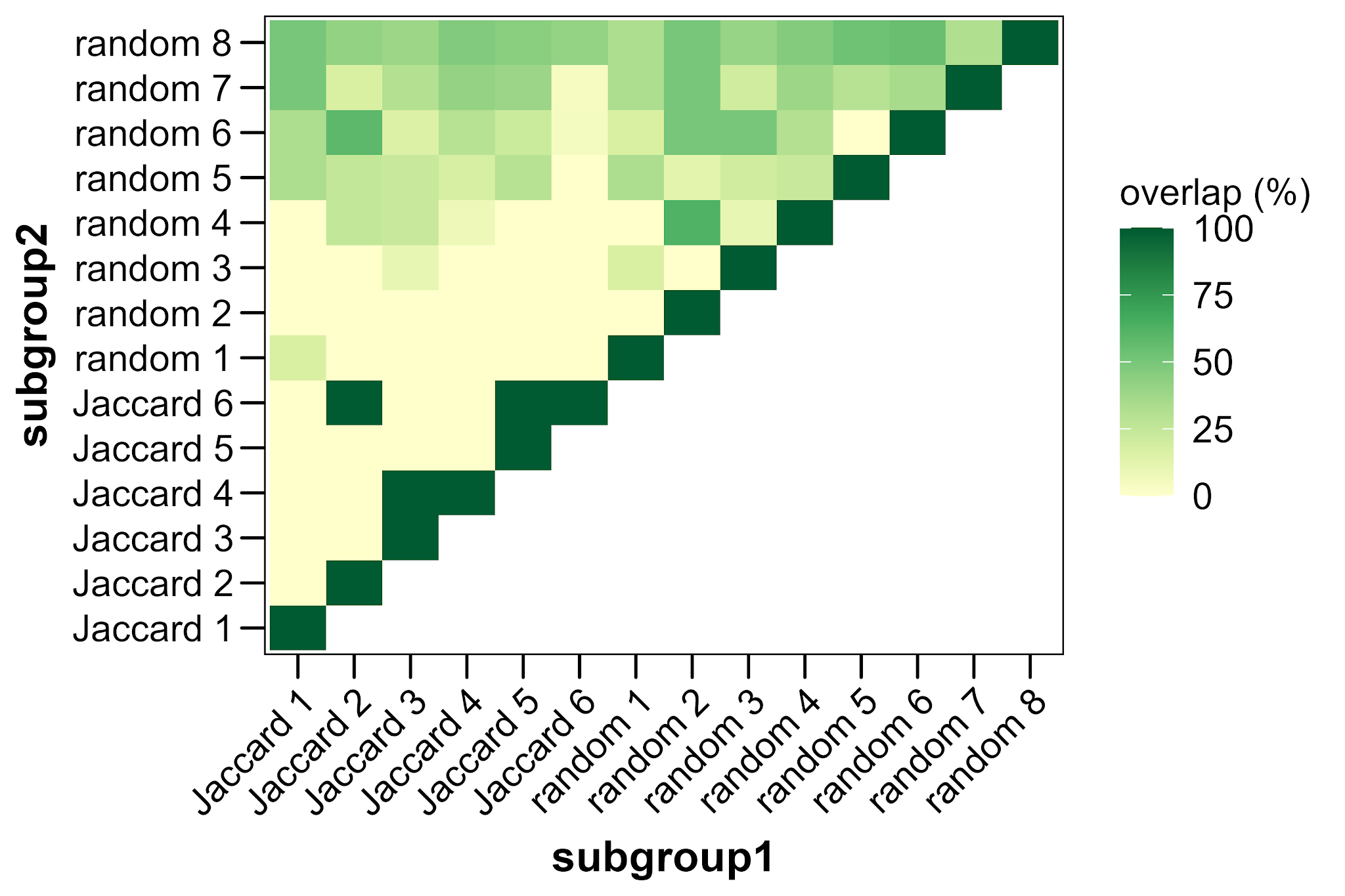

**Supplementary Figure 2 :** Overlap between subgroups of the SU2C-MARK cohort. Overlap is calculated through dividing the intersection of patients between clusters with the smallest cluster size of the two clusters compared.

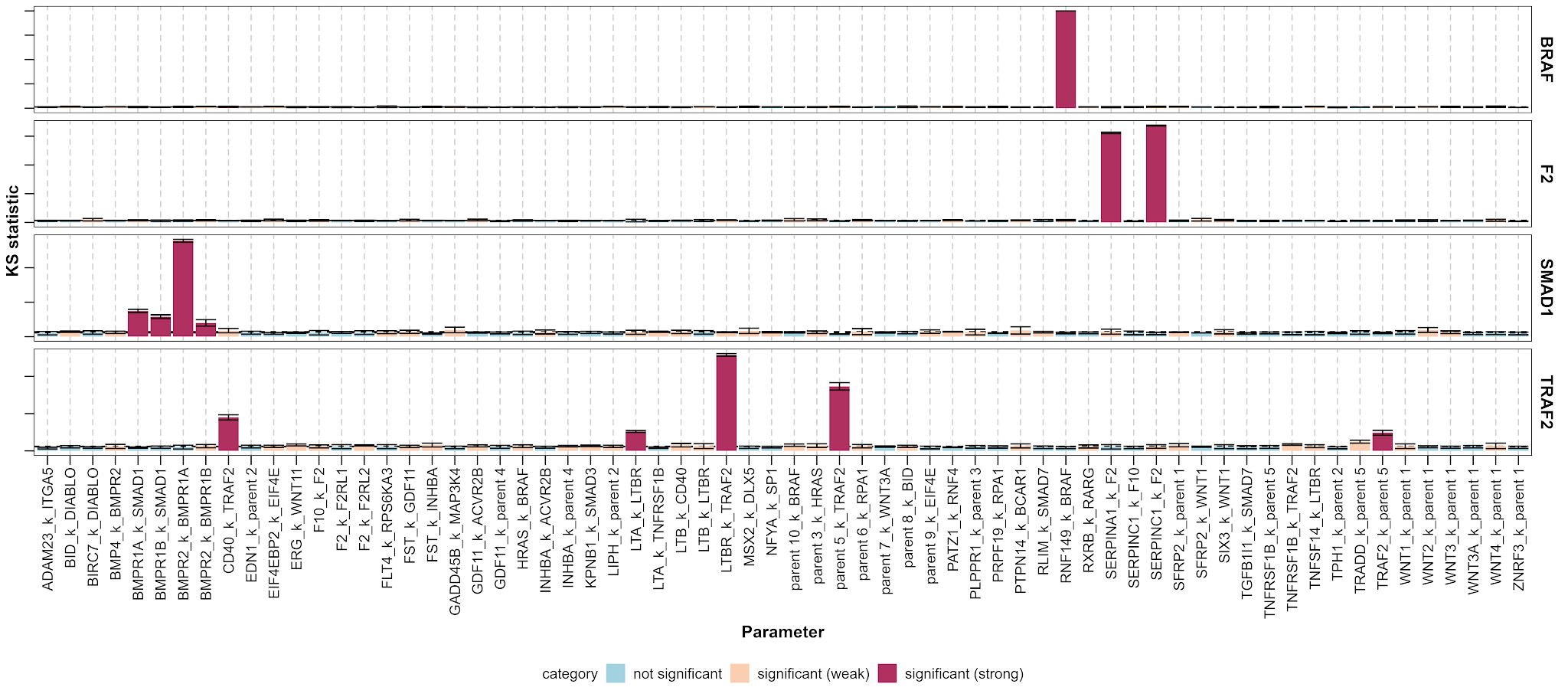

**Supplementary Figure 3 :** Results from the MPSA on the outputs BRAF, F2, SMAD1 and TRAF2. Parameters are colored based on their KS statistic, based on significance (based on threshold, which is the mean KS statistic of dummy variables). A further distinction is made between a strong (KS statistic > threshold + 3*threshold_SD) and weak significance.

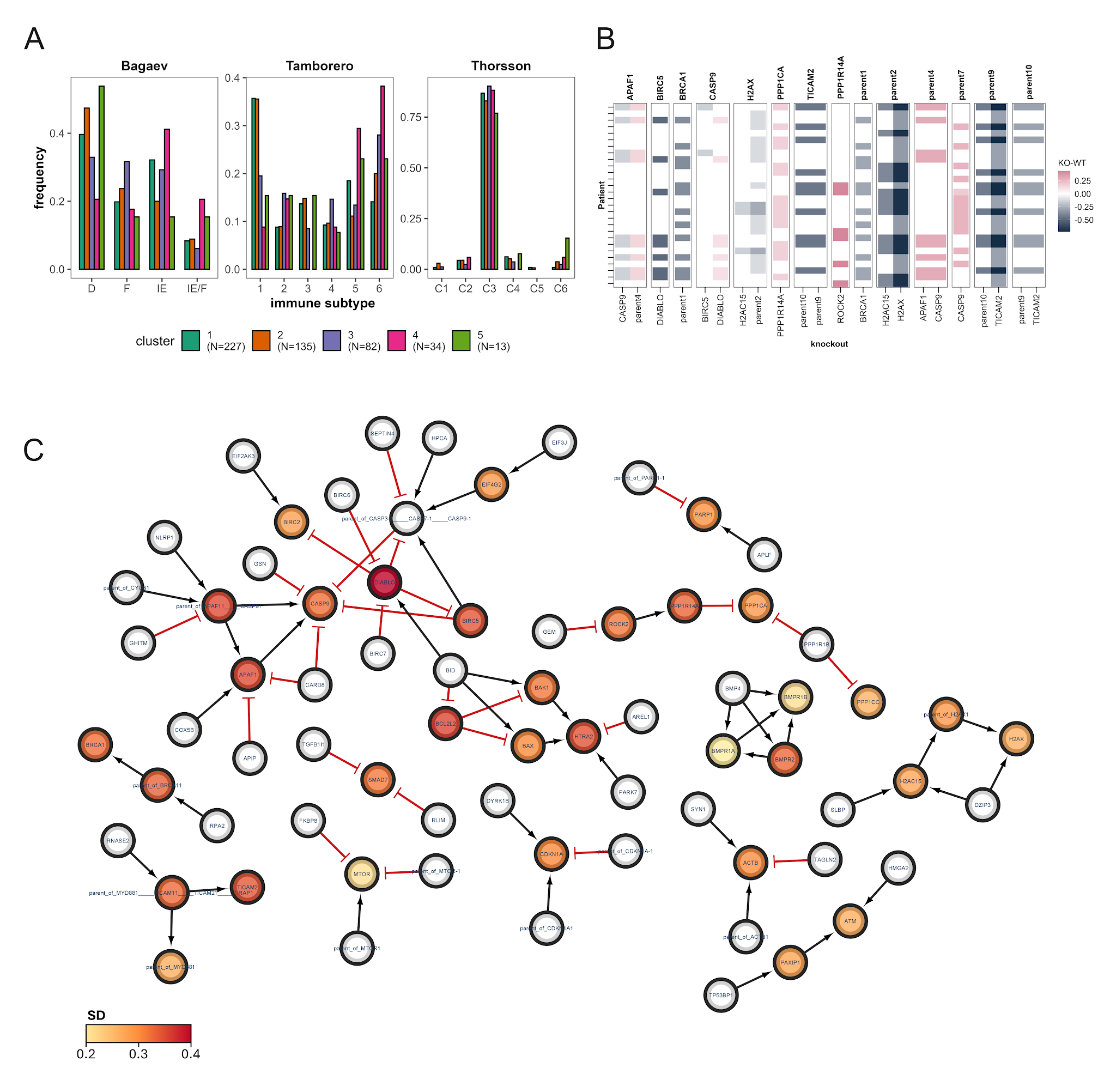

**Supplementary Figure 4** : **TCGA-KIRC cohort** **(A)**: Distribution of patients per TCGA-KIRC subgroups across immune subtypes defined by Bagaev, Tamborero and Thorsson. **(B)** In silico knockout screening of subgroup 4 of the TCGA-KIRC cohort. For every simulated protein activity, knockouts that resulted in a difference between knockout and wildtype model larger than 0.1 are visualized (KO-WT). **(C)** Standard deviation across patients in subgroup 4 of the TCGA-KIRC cohort per protein in the PKN. Only subnetworks consisting of more than two proteins are visualized.
